# A Narrow Range of Transcript-error Rates Across the Tree of Life

**DOI:** 10.1101/2023.05.02.538944

**Authors:** Weiyi Li, Stephan Baehr, Michelle Marasco, Lauren Reyes, Danielle Brister, Craig S. Pikaard, Jean-Francois Gout, Marc Vermulst, Michael Lynch

## Abstract

The expression of genomically-encoded information is not error-free. Transcript-error rates are dramatically higher than DNA-level mutation rates, and despite their transient nature, the steady-state load of such errors must impose some burden on cellular performance. However, a broad perspective on the degree to which transcript-error rates are constrained by natural selection and diverge among lineages remains to be developed. Here, we present a genome-wide analysis of transcript-error rates across the Tree of Life using a modified rolling-circle sequencing method, revealing that the range in error rates is remarkably narrow across diverse species. Transcript errors tend to be randomly distributed, with little evidence supporting local control of error rates associated with gene-expression levels. A majority of transcript errors result in missense errors if translated, and as with a fraction of nonsense transcript errors, these are underrepresented relative to random expectations, suggesting the existence of mechanisms for purging some such errors. To quantitatively understand how natural selection and random genetic drift might shape transcript-error rates across species, we present a model based on cell biology and population genetics, incorporating information on cell volume, proteome size, average degree of exposure of individual errors, and effective population size. However, while this model provides a framework for understanding the evolution of this highly conserved trait, as currently structured it explains only 20% of the variation in the data, suggesting a need for further theoretical work in this area.

---

Owing to fundamental limitations at the biophysical and biochemical levels, enzymes occasionally engage with inappropriate substrates, leading to permanent errors in the case of genome replication and transient errors in the case of transcriptional and translational products. Like all cellular traits, error rates are subject to natural selection and are expected to be persistently pushed towards low levels. However, the noise in the evolutionary process associated with random genetic drift inhibits the approach to perfection, as the response to directional selection becomes thwarted once the magnitude of incremental improvement becomes smaller than the magnitude of stochastic evolutionary forces (Ohta 1973; Hartl et al. 1985; Lynch 2007).

No example has yet been found in which natural selection has pushed the performance of a cellular feature to the level of refinement dictated by a biophysics barrier, as clearly demonstrable with genome-replication fidelity. In principle, biophysical limits at the molecular level should be independent of the cellular background. Yet, the mutation rate per nucleotide site varies ∼ 1000× across the Tree of Life, and much of this variation is consistent with patterns of variation in the power of random genetic drift (Lynch et al. 2016; Lynch and Trickovic 2020; Lynch et al. 2023), with the former scaling negatively with the effective population size and the number of genomic sites under selection. The drift-barrier hypothesis also provides a simple explanation for elevated mutation rates in somatic tissues and for the error-prone nature of DNA polymerases involved in relatively small numbers of nucleotide transactions (Lynch 2008, 2010, 2011; Lynch et al. 2016).

Because they are heritable, errors made at the DNA level can have long-term, cumulative consequences. However, nonheritable cellular errors must also impose a steady-state load on the physiological capacities of cells, even though individual error-containing molecules are transient. In this study, we conducted a comprehensive survey of species across three domains of life to characterize the rates, molecular spectra, distribution, and potential functional effects of errors in transcripts. Drawing from results on a wide phylogenetic range of species, we show that transcript-error rates are consistently orders of magnitude greater than genomic mutation rates. However, the ratio of these two error rates is generally elevated in unicellular relative to multicellular species, and as a consequence there is only a five-fold range in transcript-error rates across the Tree of Life. In an effort to understand, these relative stability of these high error rates across the Tree of Life, we present a model incorporating potential factors operating at the cellular and population-genetic levels to yield insight into how transcript-error rates might be shaped by natural selection and random genetic drift. However, the relatively weak explanatory power of the theory suggests the need to identify additional parameters to achieve a comprehensive understanding of the evolution of transcript-error rates.

## Results

### A phylogenetically diverse set of transcript-error rate estimates

To estimate the incidence of errors per nucleotide site within individual transcripts, we employed the rolling-circle sequencing method (Lou et al. 2013; Acevedo and Andino 2014), with refinements to minimize the chance of introduction of spurious errors during sample preparation and sequencing (Gout et al. 2013, 2017; Fritsch et al. 2020; Li and Lynch 2020). Briefly, the method involves isolation, fragmentation, and circularization of cellular RNAs, followed by rolling-circle reverse transcription to produce multiple linked repeats of each fragment. After high-throughput sequencing of the concatenated repeats, the individual reads of each fragment are mutually aligned, enabling the separation of true transcript errors from the substantial sporadic errors introduced into single repeats during reverse transcription and/or DNA sequencing. The ability of this method to accurately identify *bona fide* transcript errors has been thoroughly validated in a number of ways, including via evaluations of responses to damaged bases, faulty RNA polymerase, and removal of nonsense-mediated decay (Gout et al. 2013, 2017; Fritsch et al. 2020; Chung et al. 2023). Additionally, we find that the circularization procedure does not introduce bias in estimating the abundance of RNA molecules compared to regular RNAseq protocols (Figure S1).

The analyses reported here substantially expand prior work in this area by adding new data for seven bacterial species (*Caulobacter crescentus, Deinococcus radiodurans, Kineococcus radiotolerans, Mycobacterium smegmatis, Pseudomonas fluorescens, Salmonella enterica, Staphylococcus aureus*), one member of the archaea (*Haloferax volcanii*), three unicellular eukaryotes (*Chlamydomonas reinhardtii, Paramecium caudatum, Paramecium tetraurelia*), and a multicellular plant (*Arabidopsis thaliana*). In addition to evaluating the accuracies of nuclear RNA polymerases (RNAPs), those for mitochondrial and chloroplast RNAPs were examined. Combined with estimates from previous studies, this gives a broad phylogenetic perspective on transcript-error rates. One striking observation is that there is a narrow range of transcript-error rates, between 10^−6^ to 10^−5^ errors per ribonucleotide for messenger RNAs (mRNA) transcribed from nuclear/nucleoid chromosomes and RNAs from mitochondria and chloroplasts (Figure 1A). This narrow range holds true even when estimates from different types of non-coding RNAs (ncRNAs) are included (Table S1).

**Figure 1.**
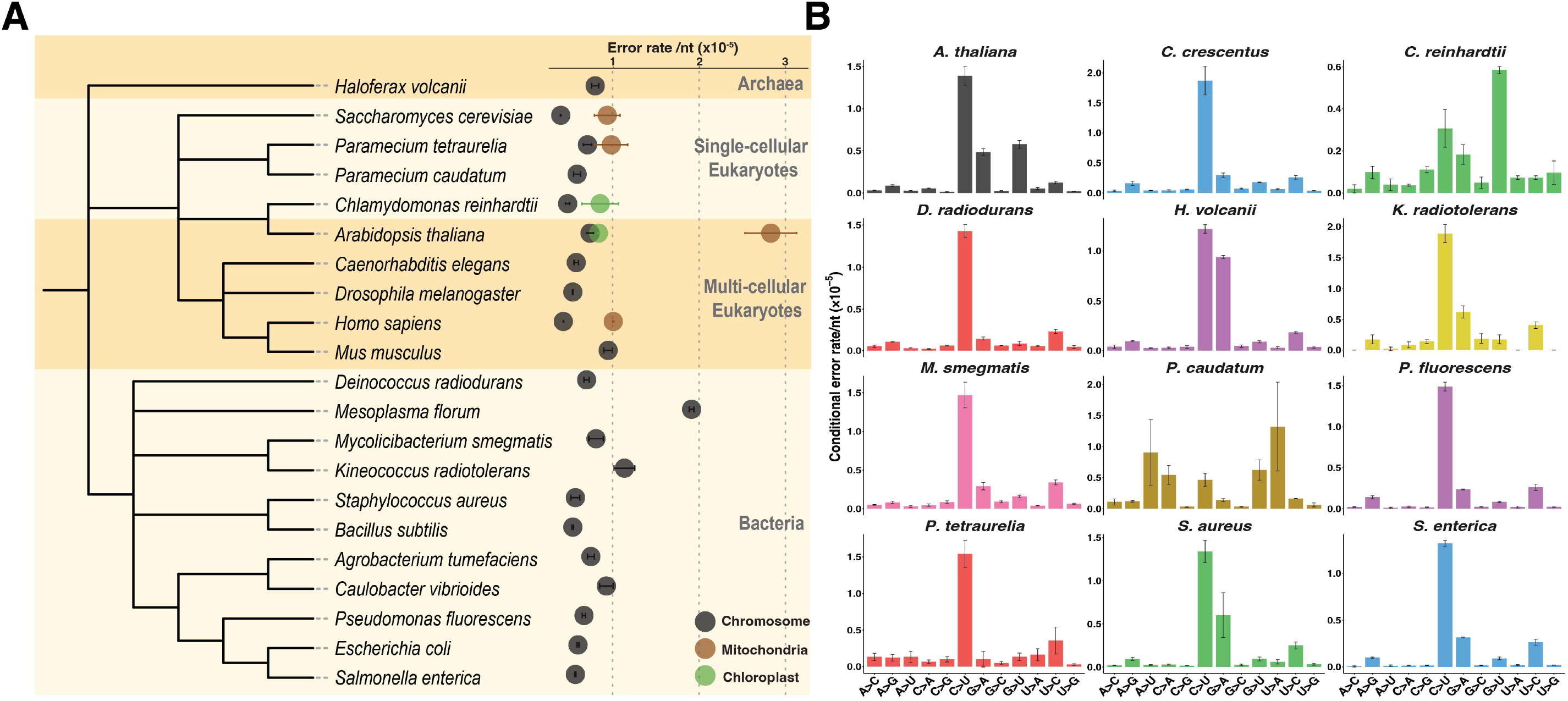
Rates and molecular spectra of transcript errors across species. **A)** A summary of all available estimates of transcript-error rates from mRNAs transcribed from nuclear/nucleoid chromosomes and RNAs from mitochondria- and chloroplast-encoded RNA polymerases. Estimates for *A. tumefaciens, B. subtilis, E. coli, M. florum, C. elegans, D. melanogaster, H. sapiens*, and *M. musculus* were obtained from previous studies using the same modified CirSeq method (Table S1). Error bars denote standard errors of the mean transcript-error rates calculated from biological replicates. The phylogeny was generated according to the NCBI taxonomy database. **B)** The molecular spectra of transcript errors from nuclear chromosomal mRNAs. The conditional error rates of each type of ribonucleotide (rNTP) substitution were calculated from the number of detected errors divided by the number of corresponding rNTPs evaluated. Error bars indicate standard errors.

Another intriguing finding is the association of the fidelities of mitochondrial and chloroplast RNA polymerases with their distinct evolutionary origins. Mitochondrial RNAP is a single-subunit enzyme resembling bacteriophage RNAP (Masters et al. 1989; Cheetham and Steitz 1999; Gaspari et al. 2004), which lacks a proofreading domain. Transcript-error rates of mitochondrial RNAPs in *S. cerevisiae, C. elegans, H. sapiens*, and *A. thaliana* are consistently on the higher end of the distribution of all observed transcript-error rates, approximately 10^−5^ errors per ribonucleotide incorporated. In contrast, chloroplast-encoded RNAP, a multi-subunit enzyme thought to originate from cyanobacterial ancestors (Börner et al. 2015), exhibits error rates similar to those of bacterial RNAPs. For example, the estimated transcript-error rates for chloroplast-encoded RNAPs in *A. thaliana* and *C. reinhardtii* are 8.3 (0.1)×10^−6^ and 8.5 (2.1)×10^−6^ errors per ribonucleotide respectively (Table S1 and S7).

### Biased molecular spectra of transcript errors

Transcriptomes consist of diverse forms of RNA molecules, including mRNAs, transfer RNAs (tRNAs), ribosomal RNAs (rRNAs), and other types of ncRNAs. In our assays, the data generated from chromosomal mRNAs were sufficiently abundant to allow cross-species comparisons for each type of ribonucleotide substitution. Conditional error rates of all 12 types of substitution were calculated by dividing the numbers of each substitution observed by the respective numbers of corresponding ribonucleotides assayed (Figure 1B). Consistent with results from previous studies (Gout et al. 2013, 2017;Li and Lynch 2020; Fritsch et al. 2020; Chung et al. 2023), the molecular spectra of transcript errors are biased towards transitions over transversions, generally dominated by C-to-U substitutions. This suggests that RNAPs have a reduced capacity for distinguishing rNTPs from the same vs. different structural categories. Two exceptions are *C. reinhardtii* and *P. caudatum*, where the spectra are dominated by G-to-U and U-to-A substitutions, respectively.

### Transcriptome-wide distributions of errors tend to be random

Previous studies suggest that transcript errors tend to be randomly distributed across genes in bacteria (Li and Lynch 2020). To further explore whether this is a general patten across a broader phylogenetic range, we evaluated the distribution of transcript errors across nuclear / nucleoid protein-coding genes. The expected number of transcript errors from each gene was calculated according to the total read coverages of the corresponding gene and the overall transcript-error rates across all chromosomal protein-coding genes. Assuming the number of transcript errors within each gene is Poisson distributed, a hotspot is identified if the observed number of errors significantly exceeds the expected number (Bonferroni-corrected *P* values of 0.05). Among the 12 species studied, 1 to 2 hotspots were detected in *K. radiotolerans, M. smegmatis, P. fluorescens, A. thaliana*, and *P. caudatum* (Table S6), consistent with the hypothesis that transcript errors are generally randomly distributed across genes in most species.

### Under-representation of potentially deleterious errors

To determine whether transcript errors are differentially distributed across functional categories of nucleotide sites, for protein-coding genes, we subdivided the latter into synonymous, missense (errors resulting in an aminoacid substitution if translated), and nonsense (errors resulting in premature stop codons). This shows that the majority of transcript errors result in missense errors, if translated (Figure 2A). To further evaluate whether errors are differentially enriched in particular functional categories, the ratios of observed to expected incidences were obtained for each error type (Figure 2B). For all species, the ratios of observed to expected rates of nonsense errors are smaller than one. The ratios for missense errors also tend to be smaller than one in almost all species, whereas synonymous errors are over-represented (as a natural consequence of the former underrepresentations). These results suggest that, in addition to cells being capable of removing some nonsense-containing transcripts (e.g., by nonsense-mediated decay), unknown mechanisms may exist for removing a fraction of missense-containing transcripts or for reducing primary error rates in the first and second positions of codons.

**Figure 2.**
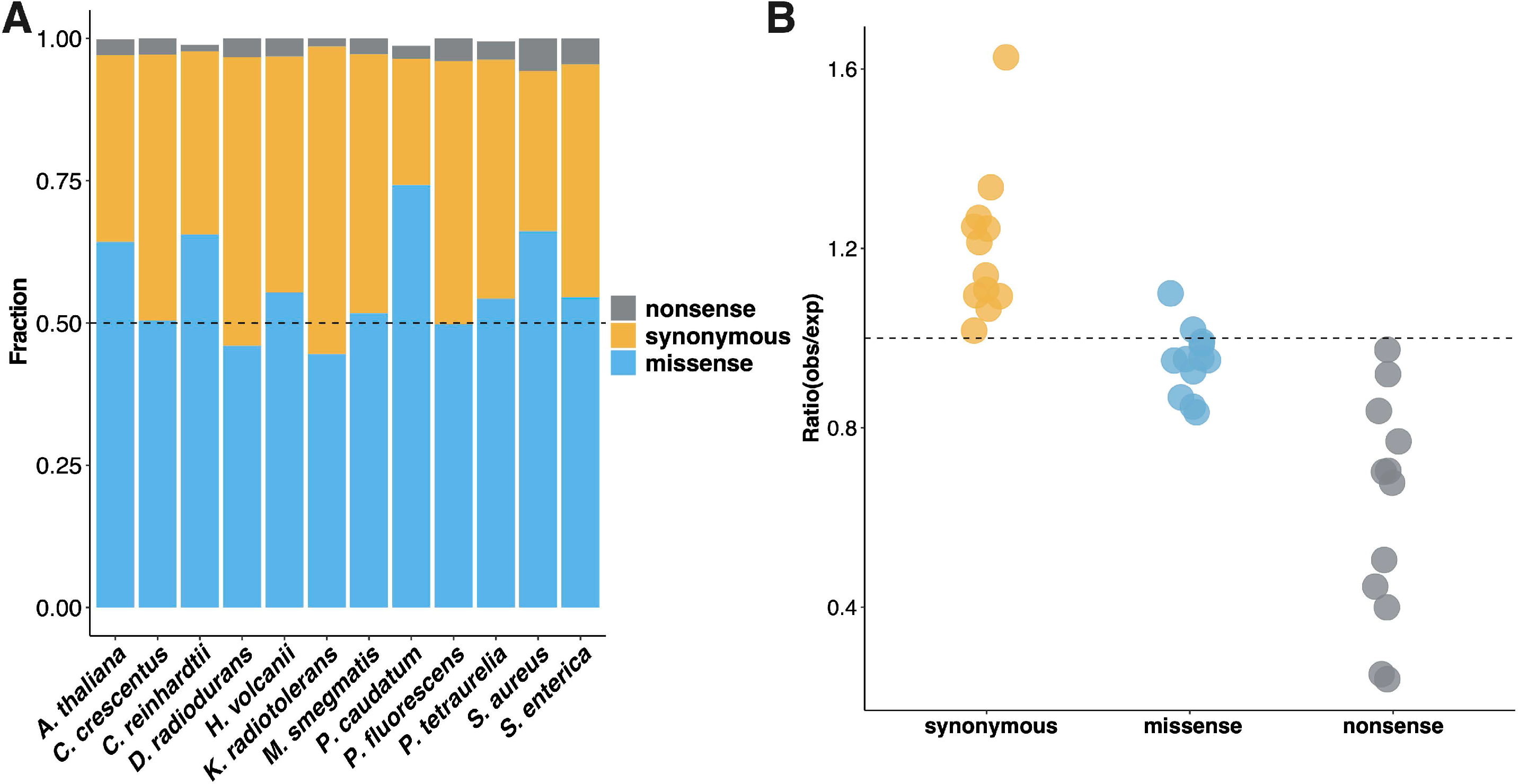
Characterization of potential functional effects of transcript errors **A)** Fractions of transcript errors from chromosomal protein-coding genes are classified as synonymous, missense, and nonsense if translated. Stop-codon loss errors detected in *A. thaliana, C. reinhardtii, P. tetraurelia*, and *P. caudatum* with minimal percentages are not displayed. **B)** The ratio of observed to expected rates for each type of errors. The expected error rates were calculated assuming a random generation of transcript errors according to the bias of ribonucleotide substitution rates and codon usages of each species, and in the absence of any correction mechanisms. Each dot represents a ratio from one species. The ratio of 1.0 is indicated by a dashed line.

### Evaluation of the variation of transcript-error rates

To obtain a broad phylogenetic perspective on the variation of transcript-error rates associated with nuclear / nucleoid protein-coding genes, in the following analyses, we include prior results for two mammals (human and mouse), a fly (*D. melanogaster*), a nematode (*C. elegans*), budding yeast (*S. cerevisiae*), and four other bacterial species (Gout et al. 2013, 2017; Fritsch et al. 2020; Li and Lynch 2020; Chung et al. 2023). We have not included results from two prior studies (Traverse and Ochman 2016; Reid-Bayliss and Loeb 2017), as these give anomalously high estimates, likely because of the more aggressive methods of mRNA fragmentation used in sample preparation (Gout et al. 2017; Meer et al. 2020).

Three striking patterns are revealed by the data (Figure 3A; Table S1). First, transcript-\ error rates are consistently orders of magnitude higher than genomic mutation rates, with the inflation being > 10^4^× in all unicellular species (except *Mesoplasma florum*), but more on the order of 10^3^× for all multicellular eukaryotes, which have elevated DNA-level mutation rates. Second, there is only a five-fold range of interspecies variation for mRNA transcript-error rates, and the range reduces to three-fold if *Mesoplasma* is excluded. Third, whereas genomic mutation rates in multicellular eukaryotes are consistently one to two orders of magnitude higher than those in unicellular species, there is no discernible difference in transcript-error rates among major phylogenetic groups. This relative constancy of the transcription-error rate across the Tree of Life is strikingly different from the orders-of-magnitude ranges of variation known for other shared traits, including metabolic rates (DeLong et al. 2010), maximum growth rates (Lynch et al. 2022), and genomic mutation rates (Lynch et al. 2016)

**Figure 3.**
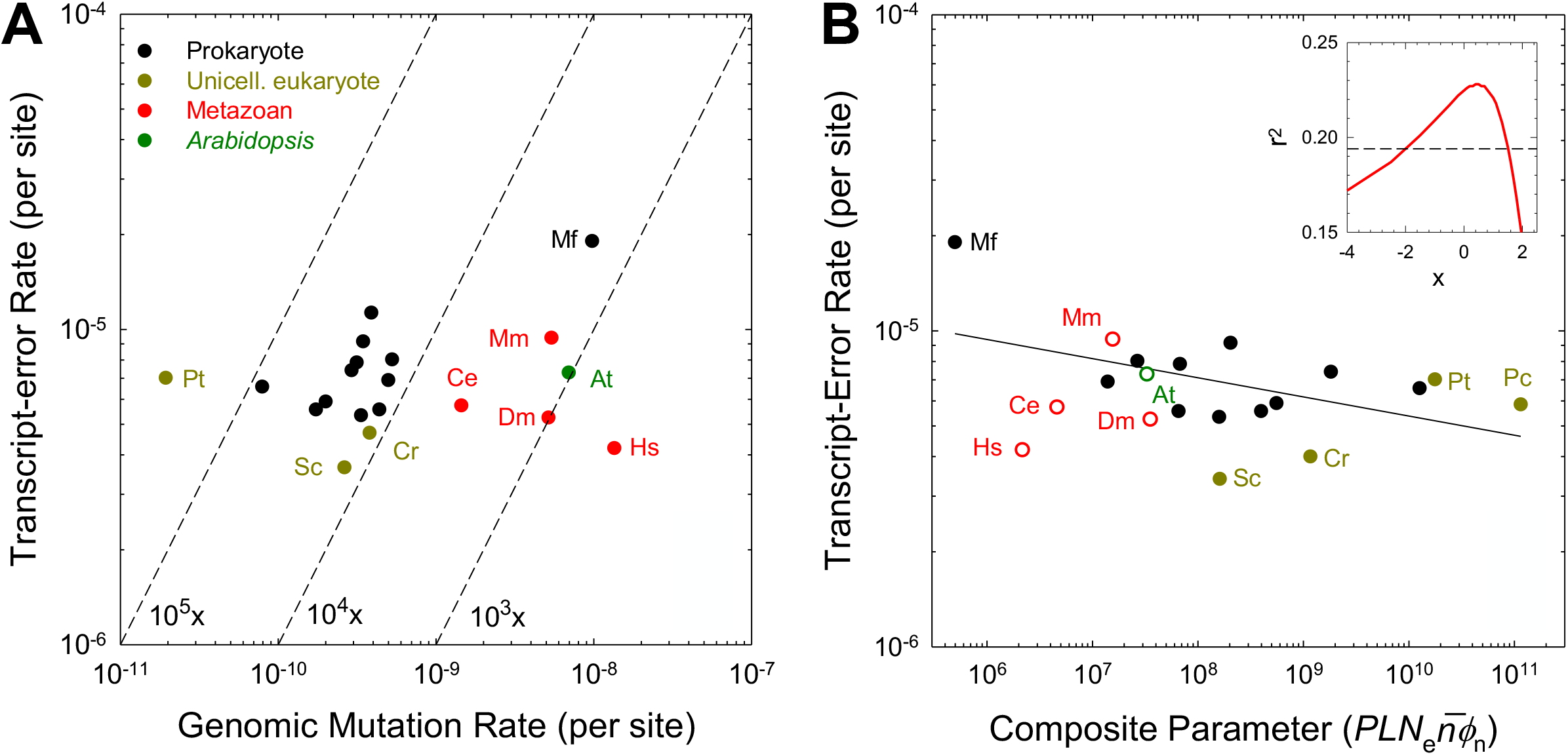
**A)** Transcript-error and genomic-mutation rates (substitutions per nucleotide site) for species across the Tree of Life. Genomic-mutation rate data are derived from Lynch et al. (2016), Long et al. (2017), and Lynch and Trickovic (2020). Diagonal dashed lines are isoclines for ratios of the two rates. **B)** Log-log regression of the observed transcript-error rates for unicellular species as a function of the composite parameter described in the text; the best fit (given by the regression line) is obtained with exponent value *x* = 0.5 (with the statistical fits with alternative *x* values given in the inset, and the support interval within which the regression remains significant at the 0.05 level denoted by the dashed line). The data points for multicellular eukaryotes are given for reference, but not used in the regression. *PL* denotes the total proteome size (summed over all codons, in megabases). Standard errors of the individual measures of transcript-error rates are generally smaller than the widths of the points. All data, including those for unlabeled bacteria (which include all species except for *Kineococcus*, for which *N*_*e*_ data were unavailable), are contained in Supplementary Table S1. *At* = *Arabidopsis thaliana; Ce* = *Caenorhabditis elegans; Cr* = *Chlamydomonas reinhardtii; Dm* = *Drosophila melanogaster; Hs* = *Homo sapiens; Mf* = *Mesoplasma florum; Mm* = *Mus musculus; Pc* = *Paramecium caudatum; Pt* = *Paramecium tetraurelia; Sc* = *Saccharomyces cerevisiae*.

Notably, these elevated rates of transcript error arise despite the capacity of proof-reading in RNA polymerases (Alic et al. 2007; Sydow and Cramer 2009) and despite the presence of multiple mechanisms of surveillance and eradication of certain classes of erroneous mRNAs (e.g., nonsense-mediated decay and nonstop decay; Graille and Séraphin 2012; Kervestin and Jacobson 2012). Nor can these high rates be explained as inadvertent by-products of elevated rates of transcription relative to DNA replication, i.e., as tradeoffs between speed and accuracy. Average speeds of transcript progression exhibit only a small range of phylogenetic variation: 46 bp/sec in *E. coli* (Golding and Cox 2004; Proshkin et al. 2010); 20 to 60 bp/sec in yeasts (Larson et al. 2012; Eser et al. 2016; Lisica et al. 2016; Ucuncuoglu et al. 2016); and 21, 23, and 56 bp/sec in *Drosophila*, rat, and human, respectively (Ardehali and Lis 2009). In contrast, replication rates are typically in the range of 100 to 1000 bp/sec in prokaryotes (Hiriyanna and Ramakrishnan 1986; Stillman 1996; Myllykallio et al. 2000), but just 10 to 50 bp/sec in yeast, flies, and mammals (summarized in Lynch 2007, 2023). Thus, transcription is much slower than replication in bacteria, whereas both processes proceed at comparable rates in eukaryotes. Finally, the low variation of transcript-error rates is inconsistent with the idea that more complex molecular machines are less error prone. The RNAPs of eukaryotes contain nearly twice the numbers of subunits per complex as those in prokaryotes (Carter and Drouin 2010), and yet the former gain nothing in terms of transcript fidelity.

### Global vs. local control of error rates

It has been suggested that in organisms with sufficiently high effective population sizes (e.g., bacteria), selection may be efficient enough to promote features that modulate error rates on a gene-by-gene basis, with such rates being adaptively reduced at genetic loci with the highest levels of expression (Meer et al. 2019). Were this to be true, genome-wide estimates of the error rate (as presented above) would not necessarily be representative of the true situation at individual nucleotide sites. Aside from the issue of whether the strength of selection on error rates at the single-gene level is ever sufficiently large to enable the evolution of local control, implicit in this argument is the assumption that errors in transcripts are not recessive to their error-free sisters. If this were not the case, lowly-expressed genes would be under stronger selection for error suppression. Although Meer et al. (2019) found evidence of a weak decline in the error rate with increasing protein abundance in *E. coli* (∼ 2-fold for the non C-to-U rate) and in yeast (∼ 1.2-fold for the C-to-U rate), there was no such pattern for total error rates (merging twelve types of base substitution-errors) and mRNA abundance in yeast (as previously pointed out in Gout et al. 2017).

To evaluate this issue more broadly, for each of the 21 species, we used the generalized linear model proposed in Meer et al. (2019) to estimate the correlation between gene-specific expression levels (mRNA abundance) and total transcript-error rates (Methods). Contrary to Meer et al. (2019), we find a significant positive correlation between error rates and expression levels in *E. coli* and four other species (and significant negative correlations for three other species), although in all cases the overall range of variation across expression level is quite small (Figure 4). Analyzing the data in other ways does not alter this conclusion (Table S4). A general adaptive hypothesis for why transcript error-rates would weakly increase with gene expression in some lineages and weakly decrease in others eludes us.

**Figure 4.**
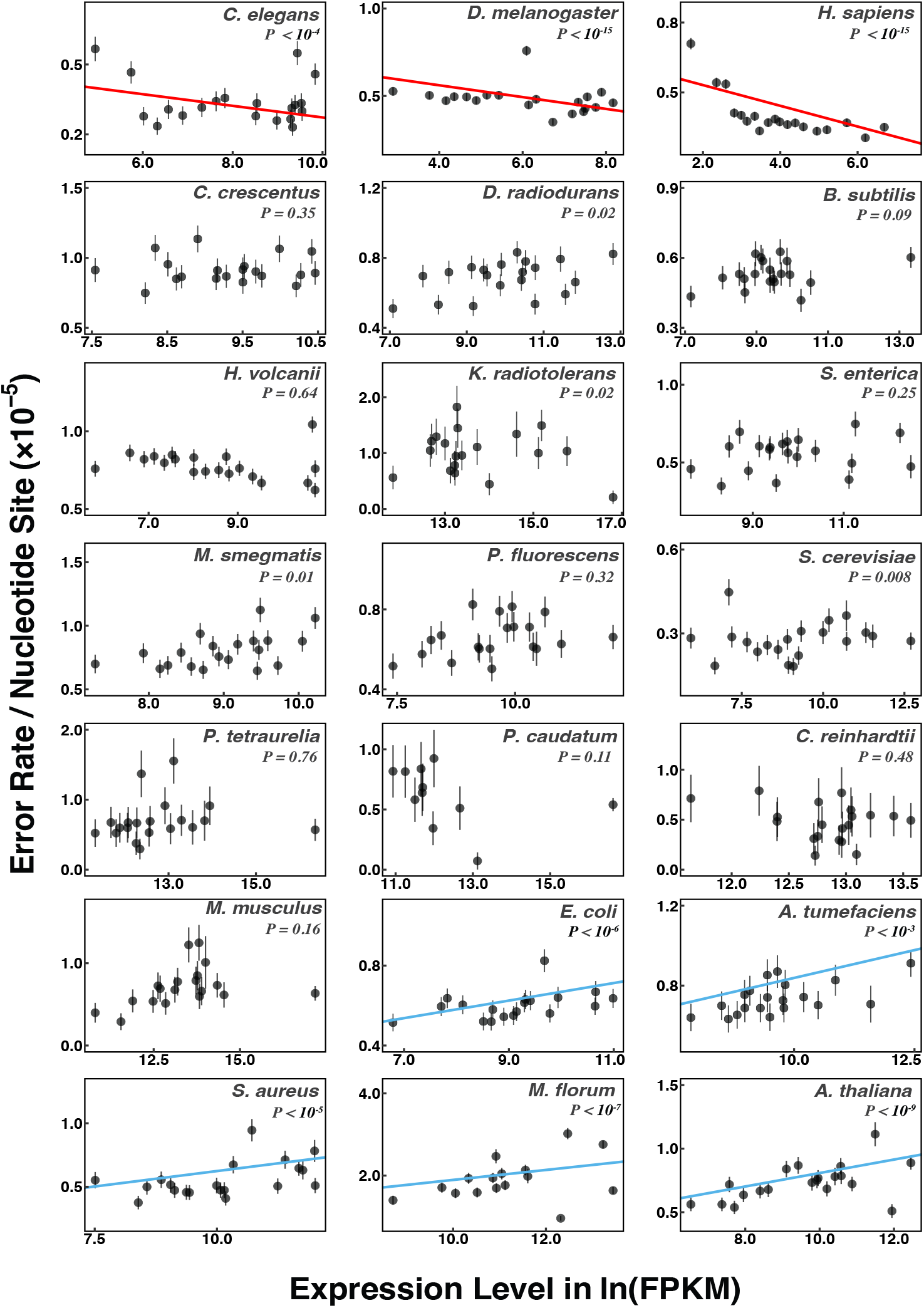
Transcript-error rates of protein-coding genes at different expression levels. A generalized linear model (Methods) was applied to expression levels (FPKM) and transcript-error rates of individual protein-coding genes to evaluate potential correlations. Expression levels of genes were obtained from CirSeq reads, which provide estimates for expression levels consistent with regular RNA-seq reads (Figure S1). Slopes with P-values less than 0.05/21 (Bonferroni correction for regression analyses for 21 species) are considered statistically significant. Significant positive and negative regressions are highlighted with blue and red lines, respectively. For visualization purposes only, protein-coding genes are ranked according to expression levels (low to high) and grouped into bins of equal cumulative width in the total number of sequenced nucleotides with 5% increments. Discrete FPKM of some genes in *M. musculus, P. caudatum*, and *M. florum* result in a few of the 20 bins being empty, and these are combined with adjacent bins on the larger FPKM side. Error bars denote standard errors of the mean error rates calculated from 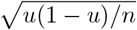, where *u* is the mean error rate of the bin and *n* denotes the number of nucleotides assayed in the corresponding bin.

The preceding analyses focus on correlations between standing levels of mRNA abundances and error rates. Might variation in rates of transcription bear on the interpretation of the observed comparative data? As noted above, progression rates of RNAP molecules engaged in transcription are quite similar among diverse organisms. There is, however, variation in the overall rates of transcript production, which is a function of the frequency of RNAP engagement.

This can be shown by noting that the steady-state number of transcripts per cell scales across species as ≃ 6700*V* ^0.4^, where *V* is the mean cell volume in *µ*m^3^ (Lynch and Marinov 2015). Given the average cell volume for a species, the size at birth is approximately 2*V*/3, so a volume equivalent to another 2*V*/3 must be produced per cell division. This implies that ∼ 5700(1 + *d*_*t*_*T*)*V* ^0.4^ transcripts must be produced per cell division, where *d*_*t*_ is the decay rate of transcripts (Supplementary Material), and *T* is the cell-division time (the term *d*_*t*_*T* accounts for transcript turnover within the life of a cell). For an *E. coli*-like cell with *V* = 1 *µ*m^3^, a cell-division time of *T* = 0.4 hours under optimal laboratory growth conditions (Lynch et al. 2022), and 4000 genes, the average rate of transcript production is then estimated to be ∼ 18 mRNAs/gene/hour. On the other hand for budding yeast *S. cerevisiae* with *V* ≃ 70 *µ*m^3^, a cell-division time of 3 hours under optimal laboratory growth conditions, and ∼ 6000 genes, the average rate of transcript production is ∼ 27/gene/hour, and assuming the same decay rate as in *S. cerevisiae*, the fission yeast *S. pombe* produces an average 32 mRNAs/gene/hour. Finally, for a *Paramecium*-like cell with *V* ≃ 250, 000 *µ*m^3^, a cell-division time of 10 hours under optimal laboratory growth conditions, and ∼ 25, 000 genes, again assuming the same decay rate as observed in yeast, the average rate of transcript production ≃ 168/gene/hour. These results show that the ∼ 10-fold range of variation in transcript production rates is not reflected in phylogenetic differences in transcript-error rates, which is inconsistent with a “wear-and-tear” hypothesis.

Taken together, the above observations suggest that transcript-error rates are not influenced by levels of transcriptional activity on either a local (gene-specific) or global (genome-wide) basis. One can further infer the difficulties in evolving gene-specific modulation of transcript-error rates in the following way. The mean number of errors per nucleotide site within a transcript across species is ≃ 7 × 10^−6^, so assuming a plausible average effect of a fully expressed mutation on fitness of 0.001 (Lynch 2024), if the error was completely dominant (i.e., there were no masking of the error effects by other transcript copies), the selective advantage of a mutant that completely eliminated errors at the site (e.g., by populating it with an unmutable nucleotide, were such a thing to exist), would be 7 × 10^−9^. If the errors were only partially dominant and/or the mutant only partially eliminated the error rate, the selective advantage would be lower (approaching 10^−9^ or smaller). Such a small advantage is beyond the reach of natural selection, given that no known species has *N*_*e*_ > 10^9^ (Lynch and Trickovic 2020). Thus, although there are small differences in the average rates of transcript-error production in genes with different expression levels in a few species, the most extreme effect (< 3×) is unlikely to be a consequence of selection as opposed to being an inadvertent by-product of other expression-associated processes (Drummond and Wilke 2009).

### Evolutionary hypotheses

The universal elevation of transcript-error rates relative to genetic mutation rates and the lack of local control of such rates strongly suggest that the strength and/or efficiency of selection for error-rate reduction is substantially reduced at the level of transcription relative to replication. The lack of patterning in Figure 3A may lead to the impression that there are no explanatory biological factors associated with interspecies variation in the transcript-error rate, and that there is a uniform optimum error rate across the Tree of Life, contrary to the situation with genomic mutation rates (Lynch et al. 2016, 2023; McCandlish and Plotkin 2016). However, the error-rate estimates displayed in Figure 3A have standard errors that are typically < 10% of the estimates, implying the presence of true (albeit small) parametric differences between species. In the spirit of motivating further thought in this area, we now consider some of the causal factors that might be involved in explaining the dispersion of the phenotypes of this unusual trait.

Adaptive hypotheses for the maintenance of error rates at an intermediate optimum by a balance between opposing selective forces embrace the common view that everything biological must have evolved to maximize organismal fitness. A specific model of this type invokes the high cost of transcription and translation fidelity resulting from the energetic cost of kinetic proofreading (Ehrenberg and Kurland 1984). The idea here is that whereas increasingly accurate transcription and translation is beneficial for cell health, too high a level of accuracy might be offset by the costs of proofreading, which generally consumes ATP/GTP and magnifies the time to progression to successful polymerization (Ehrenberg and Kurland 1984; Kurland and Ehrenberg 1984, 1987). Although such optimizing selection cannot be entirely ruled out, there are three potential issues with this hypothesis. First, it remains unclear whether the theoretical optimum level of transcript accuracy is within the reach of natural selection or beyond the drift barrier. Second, this speed vs. accuracy argument assumes that increases in accuracy can only be achieved via costly proofreading improvement rather than through improvement in accuracy of the pre-proofreading catalytic steps in transcription, which would not necessarily impose any tradeoffs. Finally, the narrow phylogenetic range in error rates, in the face of ∼ 50-fold differences in cell-division times among study species, remains unexplained, e.g., metazoans with relatively low cell-division rates do not exhibit noticeable reductions in transcript-error rates.

A radically different optimization hypothesis postulates that cellular errors have been wrongly labeled as deleterious (Peltz et al. 1999; Pezo et al. 2004; Bacher et al. 2007; Pan 2013; Ribas de Pouplana et al. 2014; Santos et al. 1999; Wang and Pan 2016). Here, the idea is that organisms “deliberately” make errors in order to expand the chemical diversity of the cell, yielding some variant molecules with large enough fitness-enhancing functions in marginal environments to offset the deleterious effects of other deviants. However, although translational (and presumably transcriptional) inaccuracies can increase during times of stress, on occasion even stochastically creating variants capable of improving a precarious situation, there is no direct evidence that error-prone processes are actively promoted and/or maintained by selection as strategies for adaptively generating variant protein pools, and plenty of evidence to the contrary (e.g., Chapter 20 in Lynch 2024). When cells are stressed, cellular functions go wrong, and there is no obvious reason why transcript-error production should be maintained as a bet-hedging strategy in healthy cells.

Multiple lines of evidence support the idea that transcript errors are commonly harmful to cell fitness. For example, knockouts of the nonsense-mediated decay pathway (which eliminates just a small subset of transcription errors containing premature stop codons) have small but discernible phenotypic effects in the yeasts *S. cerevisiae* and *S. pombe* (Leeds et al. 1992; Dahlseid et al. 1998; Mendell et al. 2000), moderate fitness effects in the nematode *C. elegans* (Hodgkin et al. 1989), and lethal effects in mice (Medghalchi et al. 2001). As many errors in mRNAs are manifest in the resultant translational products (which also have longer half lives than their mRNAs), there must be a steady-state error load carried by the many thousands of transcripts contained within individual cells, and this is reflected in proteostatic imbalance in yeast cells with elevated rates of transcription error (Vermulst et el. 2015). Again, the steady-state number of transcripts per cell ≃ 6700*V* ^0.4^, where *V* is the cell volume in *µ*m^3^ (Lynch and Marinov 2015). Approximating the transcript-error rate as 10^−5^, assuming an average transcript length of 1 kb, and accounting for transcript decay and replacement per cell lifetime (Supplemental Text), a bacterial cell with *V* = 1 *µ*m^3^ is then expected to carry a minimum ∼ 350 errors at any point in time. Eukaryotic transcripts are longer, and assuming an average of 2 kb per transcript, a yeast-sized cell with *V* ≃ 100 *µ*m^3^ would harbor ∼ 13, 600 errors (or approximately two per gene). A larger ciliate cell with *V* ≃ 10^5^ *µ*m^3^ would harbor ∼ 10^5^ errors at steady state (approximately four per gene). These numbers are relevant because although individual transcripts typically survive for only a fraction of a cell’s lifetime, their protein products can survive for the length of the cell cycle (Supplemental Text).

Given the concerns with models suggesting the adaptive maintenance of high levels of error production by stabilizing selection, we consider an alternative evolutionary model incorporating aspects of the population-genetic and cellular environment that appear most likely to influence the efficiency of selection for transcript-error reduction (Lynch 2024; Supplementary Text). Under the assumption that selection operates to minimize the incidence of transcript errors, the error-rate (*µ*^∗^) for unicellular species is expected to scale negatively with the effective population size (which increases the efficiency of natural selection), the effective proteome size (which increases the target size for error production), and the degree to which the effects of errors are manifest at the level of cellular fitness,

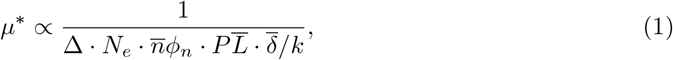

where Δ is a measure of the granularity of mutational changes in the error rates of successively improving alleles, i.e., the fractional reduction in the error rate near the drift barrier with the next improving allele; *N*_*e*_ is the effective population size; *P* is the number of protein-coding genes per genome; 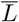 is the mean number of codons per gene; 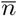 is the mean steady-state number of transcripts per cell per gene; *φ*_*n*_ is a measure of the penetrance of the effects of transcript errors (more fully developed in the following paragraph); 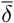 is the average reduction in fitness per transcript error if fully expressed (i.e., if manifest in all proteins associated with the gene, as in the case of a homozygous genomic mutation); and *k* is a function of the rates of decay of proteins and transcripts (defined in the Supplementary Text). As various components in the denominator of Equation 1 may be functions of *N*_*e*_, *µ*^∗^ need not strictly scale inversely with *N*_*e*_, contrary to the situation with phyogenetic variation of genomic mutation rates, which vary across taxa by several orders of magnitude (Sung et al. 2012).

The degree to which this expression extends to multicellular species remains unclear, as the possibility exists for synergistic or masking effects associated with errors contained in different cells. This general issue is even of concern within unicellular species, as specific errors will generally occur as singletons within a larger steady-state group of transcripts. The function 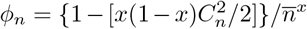 is an approximation of the average dilution factor, i.e., a measure of the average degree of expression of a transcript error, with 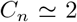 denoting the coefficient of variation in expression level among genes (Supplementary Text). When *x* = 1, 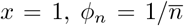, in which case the effects of individual errors are additive, and there is no net effect associated with transcript copy numbers, as the product 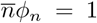. For 0 ≤ *x* < 1, the copy-number effect is more than additive, with *x* = 0 yielding *φ*_*n*_ = 1 and implying complete dominance such that 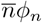 increases linearly with the number of transcripts per cell; whereas *x* < 0 implies positive synergistic effects. On the other hand, *x* > 1 implies recessivity, with larger numbers of transcripts per gene masking the effects of errors. The parameter 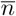 can be estimated using the scaling relationship between the mean total transcript numbers/cell and cell volume (Lynch and Marinov 2015; Supplemental Text).

Equation 1, which is certainly open to modification, illustrates that a multiplicity of factors associated with the population-genetic environment, gene structure, and the cellular environment might operate to jointly influence the overall strength of selection on transcript-error rates. Unfortunately, given that little information exists for Δ, 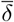, or *k*, the best that can be done currently is to consider the relationship between the transcript-error rate and the reduced composite quantity 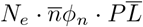, which is equivalent to the product of the effective population size and the effective number of errors per cell. This raises a potentially significant issue in that the three missing parameters may have values that scale with the parameters that are used, in which case a focus on the reduced composite parameter would be unlikely to lead to strictly inverse scaling. For this reason, we have simply performed a least-squares regression for the log-transformed transcript-error rates and species-specific estimates of 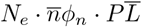 (Figure 3B).

Allowing *x* to be an additional free parameter, the optimal fit to the data for unicellular species corresponds to *x* 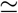 0.5 (Figure 3B), i.e., partial dominance of the effects of transcript errors. However, although the optimal fit is statistically significant (*r*^2^ = 0.23, P < 0.05 for a one-tailed test, df = 13), the support interval for *x* ranges from -2.0 (moderately synergistic) to 1.5 (moderately recessive), with more support in the direction of synergism. The slope of the optimized regression, −0.06 (SE = 0.03), is far smaller than the expectation of −1.0 if the composite parameter alone governed the evolution of the error rate (with extreme support values of *x* = −2.0 and 1.5, the best fit slopes are -0.03 and -0.07, respectively). Moreover, the marginal level of significance is eliminated entirely if the high error rate for *Mesoplasma florum* is excluded from the analysis; notably, this species also has the highest known genomic mutation rate among microbial species (Sung et al. 2012).

This weak scaling observed for transcript-error rates contrasts dramatically with the strong negative scaling known to exist for genomic mutation rates and 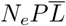 (Lynch et al. 2023). The relatively shallow level of scaling of the transcript-error rate over a 10^5^-fold range of the composite parameter may be a consequence of two effects. First, because a regression coefficient is equal to the ratio of the covariance and the variance of the explanatory variable, whenever there is sampling variance in the latter, the regression estimate will be biased towards zero. All of the components of 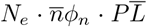 are subject to unknown levels of sampling error, but even in the extreme (and seemingly unlikely) case in which the overall variance of the composite quantity were to be inflated ten-fold by sampling error, a less biased estimate of the slope of the log-log regression would be on the order of −0.06 × 2 = −0.12, still far below the expectations for the simplest inverse-scaling model.

The second and more significant issue, alluded to above, is that the composite quantity of unknown variables 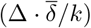 may not be independent of the remaining variables used in the analysis, 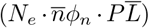. If these two composite variables were negatively correlated, the failure to include the former would lead to a reduction of the regression slope in Figure 3B, owing to the fact that the range of the latter would be greater than that of the full denominator in Equation 1. However, to achieve the inverse scaling that might be predicted by visual inspection of Equation 1, there would need to be nearly complete compensation between the variables omitted and included in the analysis. Is this plausible? The limited available data suggest that *k* may approximately double in size from small bacteria to larger eukaryotic cells (rough estimates of *k* for bacteria, yeast, and mouse are 0.16, 0.32, and 0.38, respectively; Supplementary Text). In addition, Δ is expected to decline with increasing 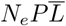, owing to the fact that as the error rate declines, there is less room for further improvement. Least clear is whether the average effects of mutations 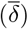 decrease with increasing 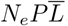. Such a pattern is unexpected from the standpoint of bioenergetic considerations (Lynch and Marinov 2015), as individual mutations imposing a bioenergetic cost must be weighed against the total energy budget of a cell, which increases isometrically with cell volume. Thus, although all of the factors in the denominator of Equation 1 would seem to be relevant to the evolution of transcript-error rates, arriving at full explanatory framework for this unusually conserved trait will require further information at the cellular level, and the necessity of including other causal factors cannot be excluded.

Finally, there are at least two reasons why the expectations of the unicellular model described above may not strictly apply to multicellular species. First, cell sizes can vary substantially among tissue types, and if fitness defects associated with errors are correlated with cell size, the use of an average cell volume to estimate the relevant mean number of transcripts per cell will be problematical. Second, there are potential higher-order effects of errors associated with multicellularity, including defects in cell adhesion, cell migration, and development, which may operate in synergistic ways that would magnify the level of selection to reduce the transcript-error rate. Nonetheless, as can be seen in Figure 3B, the error-rate estimates for multicellular species are similar to the model expectation based on unicellular taxa, with the exception of the relatively low rate for human.

## Conclusions

Our results show that, across the Tree of Life, natural selection is consistently incapable of reducing the rate of production of errors in mRNAs to the level of mutation at the DNA level, despite the fact that the polymerases involved in both kinds of activity engage with similar substrate molecules. There are at least two reasons why transcript-error rates are elevated. First, unlike heritable mutations at the DNA level which can be eradicated by selection in downstream generations, transcripts have half-lives far shorter than cell-division times so that the effects of individual errors cannot be enforced. Second, for reasons that remain unclear, translation-error rates are orders of magnitude larger than rates at the transcription level, typically in the range of 0.001 to 0.01 per codon in a wide range of experimental organisms (Lynch 2023). As a consequence, the vast majority of erroneous proteins result from errors at the translational level, and this may substantially reduce the efficiency of selection operating on transcript accuracy (Lynch and Hagner 2014; Frank 2023).

However, what remains unclear is why the transcript-error rate stalls at a level of 10^−6^ to 10^−5^ per nucleotide site and why this rate is so much more invariant across the Tree of Life than other well-studied characters such as genomic architectural features, genomic mutation rates, cellular growth rates, and swimming efficiency (Lynch 2007, 2024). The consistently high and similar rates of transcript-error production across diverse lineages do not appear to be physical by-products of high rates of polymerase transactions, which are generally lower than those for DNA-level polymerases; and even within species, there is no general tendency for error rates to increase with levels of gene expression, although a few species do exhibit a weak pattern of this sort.

In the spirit of focusing attention on potential causal variables influencing the evolution of transcript-error rates, we have introduced a model that expands on a simpler construct that has been previously presented for the evolution of genomic mutation rates, incorporating additional factors that are specifically relevant to mRNAs. However, contrary to the situation with genomic mutation rates, where there is a strong and nearly inverse relationship with *N*_*e*_ once the proteome size has been accounted for (Lynch et al. 2023), only ∼ 20% of the variation in transcript-error rates can be explained with the limited information on potentially explanatory variables. This highlights the need for additional empirical work on the fitness effects of transcript errors and how these might vary with cellular features across the Tree of Life.

## Methods

### Prokaryote strains, culture media and growth conditions

All prokaryotes were initiated from single colonies and grown in liquid culture to mid-exponential growth phase upon harvest. Details are listed in Supplementary Table S2.

### Single-celled eukaryotes

*Paramecium* strains were inherited from the lab of John Preer at Indiana University. Strain s-51 (WT *P. tetraurelia*) and strain 43c3d (WT *P. caudatum*) were maintained in wheat-grass medium inoculated with *Klebsiella* bacteria and grown at room temperature, ∼ 23°C. *Paramecium* were diluted and single cell isolates were obtained, initially growing one cell in 250 *µ*L, then expanding to 500 *µ*L, and then 1 mL in consecutive days, as cell divisions occurred. This doubling continued daily until final volumes of ∼ 5 liters were reached. 4.5 liters of *Paramecium* were then cleaned with four layers of cheesecloth laid inside a funnel, such that large debris was collected by the cheesecloth, and *Paramecium* and bacteria passed through. Next, the *Paramecium* were retained with a 15-*µ*m cloth bottomed sifting drum, through which bacteria passed. Finally, the *Paramecium* were washed and concentrated with Dryls buffer, and picked up with a plastic wide-bore Pasteur pipette, and collected in 15-mL tubes. Th latter were then spun for 8 minutes at 4000 RCF to obtain a stable pellet. After removing supernatant, the cells were resuspended following the directions of the RNEASY Midi Kit (Qiagen), and total RNA was extracted following manufacturer specifications.

Strain CC-503 cw92 mt+ of *Chlamydomonas reinhardtii*, obtained from the *Chlamydomonas* Stock Center, was grown in YA media at 20°C on a 16 hr light:8 hr cycle of ∼ 120 *µ*mol M^−2^ sec^−1^, shaken at 40 rpm. Cells were streaked on YA plates, and single colonies were picked and placed into 50-mL tubes of YA. Cells were then grown to O.D. ∼ 0.6, and harvested from liquid suspension, avoiding pelleted cells sitting at the bottom of the tube. The 49 mL of suspended cells were transferred to a new tube and spun for 1 minute at 7500 RCF. The pellet was then resuspended in 750 *µ*L of CTAB lysis buffer, and split three ways into 1.5 mL tubes, roughly 250 *µ*L per tube. RNA extraction was performed, with special care taken to lyse cell walls, using a tissue grinder in a douncing pattern to lyse the cells for 3 minutes, with breaks to disperse foaming. 150 *µ*L of 3 M sodium acetate was added to the lysate, vortexed for 10 seconds, followed by 1 mL of 100% ETOH, and vortexed again. After a 3-minute incubation on ice, the split samples were then recombined in a 15-mL tube, and spun at 7500x G for 15 minutes. The supernatant was removed, and the pellet resuspended in the contents of the Qiagen RNEASY midi kit. Total RNA was obtained and used according to the general RNA-rolling-circle protocol, as described in Gout et al. (2017).

### Plant materials

Leaf tissues of 2-week-old *Arabidopsis thaliana* Col-0 plants were ground in liquid nitrogen to a powder, and total RNA was extracted using TRIzol (ThermoFisher Scientific). rRNA was depleted by the Ribo-Zero rRNA Removal Kit (Plant Leaf) (Illumina) to enrich mRNAs for subsequent library preparations.

### Sequencing library construction

Three to four biological replicates were prepared for each species. The total RNA extraction, rRNA depletion, and construction of rolling circle sequencing library were done as previously described (Gout et al. 2017; Fritsch et al. 2020; Li and Lynch 2020). To evaluate whether circularization of RNAs during sequencing library preparations introduce bias in RNA abundances, we also prepared regular RNA sequencing libraries for *D. radiodurans, P. fluorescens, S. enterica*, and *S. aureus*. Total RNAs of these four species were split into two halves, one to construct modified CirSeq libraries and the other for regular mRNA-seq libraries. Illumina SE300 and SE100 reads were generated for CirSeq and regular RNA-seq libraries, respectively.

### Data analysis

Transcript errors were called following the analysis pipeline as previously described (Gout et al. 2017; Fritsch et al. 2020; Li and Lynch 2020) with some modifications.Different from the case of genetic mutations where all derived reads tend to show mismatches from the reference, transcript errors result in mismatches of reads at a low frequency. A statistical framework proposed in Meer et al 2020 was followed to accurately call transcript errors according to the frequency of transcript errors at individual nucleotide loci. Transcript errors were initially called if consensus base calls mismatch with the reference and the consensus-read coverage of the corresponding nucleotide locus was ≥ 50. A transcriptome-wide transcript-error rate was estimated accordingly after every nucleotide locus was surveyed. Assuming the number of transcript errors at each nucleotide locus follows a binomial distribution, the expected number of transcript errors at each locus was calculated according to the consensus-read coverage of the corresponding locus and the current estimate of the transcriptome-wide transcript-error rate. Outlier loci where the observed number of transcript errors was significantly larger than the expected one (Bonferroni-corrected *P* values of 0.05), were excluded and an updated estimate of the transcriptome-wide transcript-error rate was obtained. In the next round of iteration, revised expectations on the number of transcript errors observed from each locus were formulated by the updated estimate of transcript-error rate, outliers loci were excluded, and the estimate of the transcript-error rate was further updated. The final estimate of the transcriptome-wide transcript-error rate was obtained after convergence.

To evaluate gene expression levels, the Tophat-Cufflink pipeline was followed to calculate fpkm of each gene. To evaluate whether bias on the estimation of RNA abundances were introduced by circularization, only CirSeq reads that contain tandem repeats were used for the expression level analysis.

### Estimates of effective population sizes

The effective population sizes (*N*_*e*_) of prokaryotes in this study (not previously reported) were estimated from measures of silent-site heterozygosity in protein-coding genes (*π*_*s*_), which has an expectation of 2*N*_*e*_*µ*_*g*_, where *µ*_*g*_ denotes the base-substitution rate per nucleotide per generation, available from previous mutation-accumulation studies (Supplementary Table 1). *π*_*s*_ was estimated from nucleotide heterozygosity of four-fold degenerative sites across the whole genome from data deposited at NCBI (Supplementary Table S3), and the data used herein are drawn from compilations in Lynch et al. (2023).

### Generalized linear model for relating expression levels and transcript-error rates

To evaluate the potential correlation between expression levels and transcript-error rates, a generalized linear model was constructed, following the earlier procedures of Meer et al. (2020),

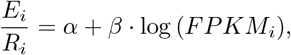

where *E*_*i*_ is the number of transcript errors at locus *i, R*_*i*_ is the number of ribonucleotides assayed at the locus, and *FPKM*_*i*_ is the gene expression level. To better evaluate this linear model considering sampling errors, we can multiply *R*_*i*_ on both sides and evaluate the following model,

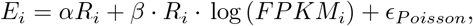

where the final term denotes the residual error. P-values of the slope (*β*) were obtained by performing ANOVA tests with the *χ*^2^ option in R to evaluate whether including the slope significantly improves the generalized linear model.

## Data availability

Sequencing data for prokaryotes, *A. thaliana, C. reinhardtii, P. caudatum*, and *P. tetraurelia* are deposited in NCBI with the BioProject Number PRJNA778257, PRJNA791609, PRJNA847594, PRJNA844889, and PRJNA839092, respectively.

## Code availability

Source code for analysis is available at https://github.com/LynchLab/CirSeq4TranscriptErrors.

## Acknowledgments

This research was supported by the Multidisciplinary University Research Initiative awards W911NF-09-1-0444 and W911NF-09-1-0444 from the US Army Research Office, National Institutes of Health award R35-GM122566-01, National Science Foundation award DBI-2119963, and Grant 735927 from the Moore-Simons Project on the Origin of the Eukaryotic Cell to ML, National Institutes of Health award GM077590 and Howard Hughes Medical Institute Investigator funds to CSP, and National Institutes of Aging award R01-AG054641-01 to MV.

