## Supplementary text and figure for "A Narrow Range of Transcript-error Rates Across the Tree of Life"

### Theory for the Evolutionary Bounds on the Transcript-error Rate

Based on numerous direct observations on the effects of random amino-acid substitutions on protein performance (Keightley and Eyre-Walker 2007; Huber et al. 2015; Kim et al. 2017; Lynch et al. 2017; Booker and Keightley 2018), as well as results from numerous large-scale population-genetic analyses (Axe 2000; Guo et al. 2004; Lind et al. 2010; Roscoe et al. 2013; Firnberg et al. 2014), we start with the assumption that the majority of errors in transcripts leading to nonsynonomous changes are deleterious upon translation. We wish then to evaluate the expected selective advantage of a genomic variant that improves transcriptional fidelity (or conversely the disadvantage of a variant that exacerbates the transcript-error rate), with the ultimate goal of ascertaining the degree to which such variants have substantial enough effects relative to the power of genetic drift to be perceived by selection. To achieve such an understanding, several factors must be considered: 1) the expected number of errors per transcript that manifest at the amino-acid sequence level; 2) the total steady-state number of transcripts/cell associated with each gene; and 3) the fitness effects of such errors.

As we assume that the fitness effects of transcript errors are revealed after translation, the following derivations will consider the rate of nonsynonymous errors per codon in mRNAs, $u$. Letting the error rate per nucleotide site be $\mu$, because there are 3 nucleotide sites per codon, and $\sim 3/4$s of nucleotide substitutions cause an amino-acid substitution (as a consequence of the structure of the genetic code), $u \simeq 9\mu/4$. Letting $L_i$ be the number of amino acids (codons) in a protein of type $i$, the numbers $(j)$ of erroneous amino acids in individual protein molecules of this type will be approximately Poisson distributed with expectation $uL_i$,

$$P(j|uL_i) = \frac{e^{-uL_i}(uL_i)^j}{j!}. \tag{S1}$$

Because transcript errors are generally singular events, and the mean number of protein molecules per active gene is generally $\gg 1$, individual variant proteins within a cell will generally be just a fraction of the total pool of molecules for specific genetic loci. This raises the question of the degree to which the fitness effects of single transcript errors are manifest at the cellular level. As with variant alleles at a genetic locus, transcription and translation errors might behave in an additive, recessive, or dominant fashion, with the magnitude of the latter two conditions depending on the number of transcripts per cell. Thus, there is a need to know the fraction of the pool of proteins present in a cell at any one time that are derived from a particular transcript (which might contain a unique transcript error).

Letting $\widetilde{n}_t$ be the steady-state number of transcripts per cell for a particular gene, the total number of transcripts produced per cell cycle is

$$n_t = \widetilde{n}_t \left(1 + \frac{d_t T}{\ln(2)}\right), \tag{S2a}$$

where $d_t$ is the transcript decay rate, and $T$ is the cell-division time. Likewise, the total number of proteins produced per cell cycle is

$$n_p = \widetilde{n}_p \left(1 + \frac{d_p T}{\ln(2)}\right). \tag{S2b}$$

It then follows that the fraction of proteins in the cell associated with one particular transcript at any time is approximately

$$\frac{1}{\widetilde{n}_t} \left(\frac{\ln(2) + d_p T}{\ln(2) + d_t T}\right) = \frac{k}{\widetilde{n}_t}. \tag{S3}$$

The fact that decay rates of transcripts are generally larger than those for proteins leads to the expectation that $k < 1$, although rough estimates can only be obtained for a few organisms. For *E. coli* and a few other bacteria, $d_t \simeq 10$/hour (Bernstein et al. 2002; Hambraeus et al. 2003; Taniguchi et al. 2010; Dressaire et al. 2013), whereas average $d_p \simeq 0.2$/hour for several bacteria (Lahtvee et al. 2014; Trötschel et al. 2013), so assuming $T \simeq 0.4$ hours for a bacterial cell growing at maximum rate, $k \simeq 0.16$. Similarly, for the yeast *S. cerevisiae,* $d_t \simeq 5$/hour (Wang et al. 2002; Neymotin et al. 2014), $d_p \simeq 1.4$/hour (Belle et al. 2006), and assuming $T \simeq 3$ hours at maximum growth rate, $k \simeq 0.32$. Finally, for mouse fibroblast cells, $d_t \simeq 0.1$/hour and $d_p \simeq 0.02$/hour (Schwanhäusser et al. 2011), which under the assumption of $T \simeq 24$ hours at maximum growth rate leads to $k \simeq 0.38$. These limited observations suggest that $k$ may be relatively constant across species, in which case its exact value will be irrelevant to the scaling relationships determined below, and it will be ignored for the time being, assuming relative constancy across species and genes.

Thus, given that the average fraction of protein molecules for a particular locus associated with individual transcripts is inversely proportional to the average transcript number, letting

$\widetilde{n}_{t,i}$ be the number of steady-state transcripts per cell for protein $i$, a flexible function that allows for alternative modes of dilution of effects is

$$f(\widetilde{n}_{t,i}) = \frac{1}{\widetilde{n}_{t,i}^x}, \tag{S4}$$

which equals 1.0 when $\widetilde{n}_{t,i} = 1$ (effects are fully felt), and converges to 0.0 (effects are completely masked) as $\widetilde{n}_{t,i} \to \infty$ at a rate that depends on the exponent $x$. When $x = 1$, $f(\widetilde{n}_{t,i}) = 1/\widetilde{n}_{t,i}$, and the number of copies of a protein has no effect, as the number of error-containing proteins, which is proportional to $\widetilde{n}_{t,i}$, is compensated by the dilution effect, i.e., $\widetilde{n}_{t,i} \cdot f(\widetilde{n}_{t,i}) = 1$. Values of $0 < x < 1$ result in a relatively slow decline in the dilution effect with increasing $\widetilde{n}_{t,i}$ (with $x = 0$ implying complete dominance), i.e., synergism of errors, whereas $x > 1$ results in a relatively rapid decline in $f(\widetilde{n}_{t,i})$ (i.e., increasingly recessive effects of errors). More complicated two-parameter models that allow for non-log-linear behavior might be warranted, but direct empirical observations will be required to reveal the utility of such nuances.

For each locus $i$, the expected fractional reduction in fitness associated with the transcript-error burden ($s_i$) will depend on the number of transcripts per cell over which the errors are distributed $\widetilde{n}_{t,i}$, the degree of expression of individual errors $f(\widetilde{n}_{t,i})$, the distribution of the numbers of errors per protein $P(j|uL_i)$, and the average reduction in fitness associated with fully expressed deleterious mutations. From the references cited above, it is known that the latter is generally $< 0.1$, and based on the transcript-error rates reported herein and typical gene lengths, the number of errors per protein will generally be $\ll 10$. Thus, letting $\delta$ be the fitness loss per single error in a single transcript if fully revealed, and assuming independent effects of individual errors, the total reduction in fitness resulting from a protein containing $j$ errors is $(1-\delta)^j \simeq 1 - e^{-j\delta}$. It then follows that

$$s_i = \widetilde{n}_{t,i} \cdot f(\widetilde{n}_{t,i}) \cdot \sum_{j=1}^{L_i} P(j|uL_i) \cdot (1 - e^{-j\delta_i}), \tag{S5a}$$

$$\simeq \widetilde{n}_{t,i} \cdot f(\widetilde{n}_{t,i}) \cdot \left[1 - \exp\left(-\frac{uL_i\delta_i}{1+\delta_i}\right)\right], \tag{S5b}$$

$$\simeq \widetilde{n}_{t,i} \cdot f(\widetilde{n}_{t,i}) \cdot (uL_i) \cdot \delta_i, \tag{S5c}$$

where the latter approximation assumes $\delta_i \ll 1$.

Further noting that the error loads at each target locus will be $\ll 1$, and assuming independent effects across loci, relative to a baseline fitness of 1.0 for an error-free individual, the selective disadvantage of a genotype with error rate $u$ is

$$s(u) \simeq 1 - \exp\left(-\sum_{i=1}^{P} s_i\right), \tag{S6}$$

where $P$ is the number of loci in the proteome. Recalling Equation S5c, the selection differential between two variants with transcript-error rates $u$ and $(1+\Delta)u$ is then

$$s(\Delta) \simeq \sum_{i=1}^{P} \widetilde{n}_{t,i} \cdot f(\widetilde{n}_{t,i}) \cdot \Delta \cdot u \cdot L_i \cdot \delta_i, \tag{S7}$$

where transformation to the additive scale follows from the fact that the terms in the exponent are $\ll 1$. Further simplification is possible if it is assumed that the copy number effects, $\widetilde{n}_{t,i} \cdot f(\widetilde{n}_{t,i})$, are independent of the lengths of loci, $L_i$, and error effects, $\delta_i$,

$$s(\Delta) \simeq [\,\overline{n} \cdot \phi_n\,] \cdot [\,\Delta \cdot u \cdot P\overline{L}\,] \cdot \overline{\delta}, \tag{S8}$$

where overlines denote mean values, and $\phi_n = \{1-[x(1-x)C_n^2/2]\}/\overline{n}^x$, obtained by Taylor-series expansion of $\widetilde{n}_{t,i} \cdot f(\widetilde{n}_{t,i})$, is the average dilution factor, with $\overline{n}$ and $C_n$ denoting the mean and coefficient of variation in expression level.

Equation S8 shows that the genome-wide fitness consequences of transcript errors are a function of three quantities: 1) the net influence of the steady-state numbers of transcripts per gene and the degree of dilution of error effects (the first bracketed expression); 2) the total error rate per expressed proteome, $uP\overline{L}$; and 3) the average effect of an undiluted amino-acid substitution, $\overline{\delta}$, which must be equivalent to the average homozygous effect of a genomic mutation. Our ultimate goal is to define the drift barrier to the evolution of transcript fidelity ($\mu^*$), i.e., the error rate at which the next incremental improvement $s(\Delta) \simeq 1/N_e$,

$$\mu^* \propto \frac{1}{\Delta \cdot N_e \cdot \overline{n}\phi_n \cdot P\overline{L} \cdot \overline{\delta}}. \tag{S9}$$

The granularity of mutational changes in alleles influencing the error rate ($\Delta$) operates as a simple scaling factor, but does not change the form of the scaling relationship with respect to other cellular features – the higher the value of $\Delta$, the greater the difference of allelic effects, and hence the greater the efficiency of selection for a lower error rate.

These results show that the scaling properties of the drift barrier depend on both the population-genetic environment ($N_e$) and the cellular environment (the remaining terms in Equation S9). As these two sets of features are not necessarily independent, a strict inverse relationship between the evolved error rate and $N_e$ may not hold. To gain further insight into these mutual dependencies, we consider the ways in which some of the terms in Equation S9 scale with organism size. From Lynch and Trickovic (2020),

$$N_e \simeq (8 \times 10^7)V^{-0.18}, \tag{S10}$$

where a conversion from dry weight (in the original paper) to organism volume ($V$ in $\mu$m$^3$) has been made using a universal relationship of dry weight (ng) $= 0.00057V^{0.92}$ (Lynch 2023). From Lynch and Marinov (2015),

$$\overline{n} \simeq 3V^{0.28}, \tag{S11}$$

and for the dilution-factor effect, $C_n = 2$ can be applied as a first-order approximation (based on an absence of phylogenetic pattern $C_n$ and an observed range of 1 to 3).

**Supplementary Tables and Figure**

**Table S1.** Summary of results, estimates, and citations used in analysis and hypothesis testing.

**Table S2.** Prokaryote strains, culture media, and growth conditions.

**Table S3.** Estimates of effective population sizes for prokaryotic species.

**Table S4.** Details of regression analyses of transcript-error rates vs. expression levels for protein-coding genes. Expression levels of genes (FPKM) were obtained from CirSeq reads, which provide estimates for expression levels consistent with regular RNA-seq reads (Figure S1). In Method 1, the generalized linear model from Meer et al. (2019) was adapted (Methods). This Poisson distribution-based model was applied to individual protein-coding genes to evaluate potential correlations between expression levels and transcript-error rates. P-values of the slope were obtained by performing ANOVA tests with the $\chi^2$ option in R to evaluate whether including the slope significantly improves the generalized linear model. In Method 2, least-squares linear regression analyses were carried out on transcript-error rates and log-transformed expression levels of individual protein-coding genes. In Method 3, the only difference from Method 2 is that expression levels of individual genes were not log-transformed, i.e., the least-squares linear regressions were carried out on arithmetic-arithmetic scales. Breusch-Pagan tests were used to test for homoscedasticity in linear-regression models generated by Method 2 and 3 (Table S5). In Method 4, logistic regression analyses were performed on transcript-error rates and expression levels of individual protein-coding genes on arithmetic-arithmetic scales. Regressions with P-values $< 0.05/21$ (Bonferroni correction for regression analyses for 21 species) are considered statistically significant. Significant positive and negative regressions are highlighted in blue and red respectively.

**Table S5.** Checking homoscedasticity in linear-regression models using the Breusch-Pagan test. Homoscedasticity of linear-regression model fits relating transcript-error rates with gene-expression levels was assessed using Breusch-Pagan tests. Two scales were used for constructing linear-regression models. On the logarithmic-arithmetic scale (Method 2 in Table S4), gene-expression levels were log-transformed before constructing linear-regression models. On the arithmetic-arithmetic scale (Method 3 in Table. S4), gene-expression levels and transcript-error rates were used directly for regression models.

**Table S6.** Hotspot genes for transcript errors detected in each species.

**Table S7.** Protein-coding genes used to estimate transcript-error rates of chloroplast-encoded RNAPs.

**Figure S1.** Least-square regressions of gene expression levels estimated from regular RNA-seq and modified CirSeq. Constructions of the modified CirSeq library requires a circularization procedure of RNA molecules that might result in a bias in estimates of expression level of different genes. To evaluate this issue, we prepared regular RNA-seq and modified CirSeq libraries in parallel for four species. FPKM (Fragments Per Kilobase of transcript per Million mapped reads) calculated from two types of libraries are positively correlated, indicating procedures of preparing the modified CirSeq library do not introduce biases in abundances of RNA fragments derived from different genes. Standard errors of the regression coefficients are given in parentheses.

Supplementary Figure 1

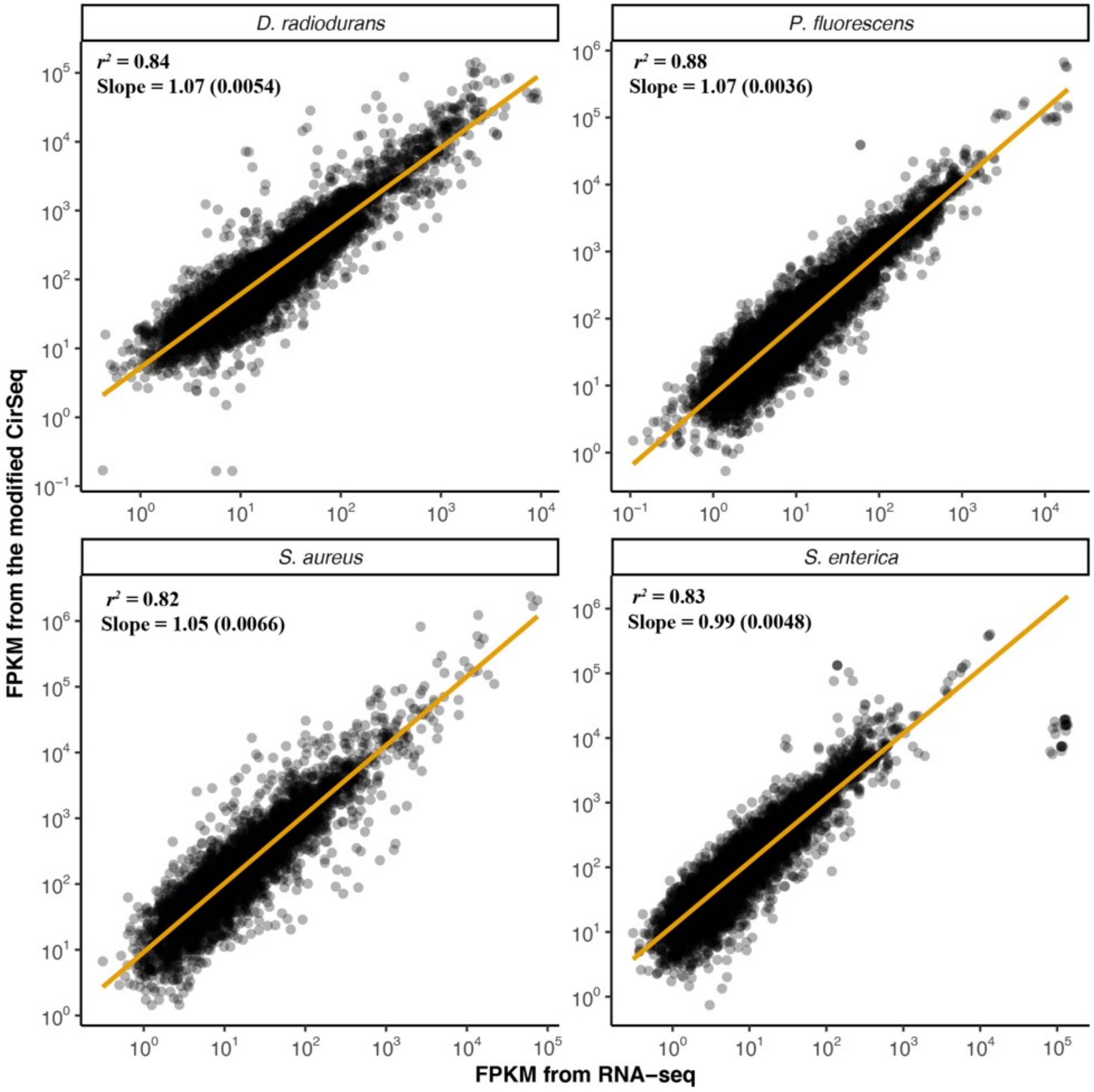
